# FBXL4 deficiency promotes mitophagy by elevating NIX

**DOI:** 10.1101/2022.10.11.511735

**Authors:** Hannah Elcocks, Ailbhe J. Brazel, Katy R. McCarron, Manuel Kaulich, Koraljka Husnjak, Heather Mortiboys, Michael J. Clague, Sylvie Urbé

**Affiliations:** Molecular Physiology and Cell Signalling, Institute of Systems, Molecular and Integrative Biology, University of Liverpool, Liverpool, L69 3BX, UK; Department of Biology, Maynooth University, Maynooth W23 F2K6, Ireland; Institute of Biochemistry II, Goethe University, Medical Faculty, University Hospital, 60590, Frankfurt am Main, Germany; Frankfurt Cancer Institute, 60596, Frankfurt am Main, Germany; Sheffield Institute for Translational Neuroscience (SITraN), University of Sheffield, Sheffield, UK

**Keywords:** FBXL4, BNIP3, NIX, Mitochondria, Mitophagy, E3 ligase, Ubiquitin, CRISPR screen

## Abstract

The selective autophagy of mitochondria is linked to mitochondrial quality control and is critical to a healthy organism. We have conducted a CRISPR/Cas9 screen of human E3 ubiquitin ligases for influence on mitophagy under both basal cell culture conditions and following acute mitochondrial depolarisation. We identify two Cullin RING ligases, VHL and FBXL4 as the most profound negative regulators of basal mitophagy. We show that these converge through control of the mitophagy adaptors BNIP3 and BNIP3L/NIX through different mechanisms. FBXL4 suppression of BNIP3 and NIX levels is mediated via direct interaction and protein destabilisation rather than suppression of HIF1α-mediated transcription. Depletion of NIX but not BNIP3 is sufficient to restore mitophagy levels. Our study enables a full understanding of the aetiology of early onset mitochondrial encephalomyopathy that is supported by analysis of a disease associated mutation. We further show that the compound MLN4924, which globally interferes with Cullin RING ligase activity, is a strong inducer of mitophagy providing a research tool in this context and a candidate therapeutic agent for conditions linked to mitochondrial dysfunction.

## Introduction

The selective disposal of damaged mitochondria by autophagy, also known as mitophagy, is a critical element of mitochondrial quality control. Defects in mitophagy have been linked to neurodegenerative ailments such as Parkinson’s and Alzheimer’s Disease and many other pathophysiological conditions (Fritsch *et al*, 2019; Rodolfo *et al*, 2018; Sorrentino *et al*, 2017). In Parkinson’s Disease, some patients show loss of function in PINK1 or PRKN genes, which render cells unable to clear damaged mitochondria (Agarwal & Muqit, 2022). PRKN encodes an RBR E3 ligase called Parkin, which is itself activated by PINK1 following damage to mitochondria. Parkin and ubiquitin are both direct substrates of PINK1, and pSer65-ubiquitin (pUb) is required for recruitment and full activation of Parkin, leading to a rapid feed-forward amplification loop and extensive ubiquitylation of mitochondrial outer membrane proteins (Bingol & Sheng, 2016; Harper *et al*, 2018; Pickles *et al*, 2018). This ubiquitin coat serves to recruit ubiquitin binding domain encoding adaptors (ubiquitin receptors) that link the mitochondrial and autophagosomal membranes, via LC3 interacting regions (LIR) (Lazarou *et al*, 2015). However, in organisms such as mice and flies, the majority of constitutive or basal mitophagy is independent of PINK1 and Parkin (Lee *et al*, 2018; McWilliams *et al*, 2018).

Based on our past studies of PINK1 and Parkin-dependent mitophagy, we have proposed that other ubiquitin E3 ligases can prime a seed population of mitochondrial proteins with ubiquitin, which can then be phosphorylated by PINK1 to unleash the Parkin-mediated ubiquitylation cascade (Marcassa *et al*, 2018; Rusilowicz-Jones *et al*, 2020). Importantly, in cell lines that do not express Parkin, we have observed a PINK1-dependent component of mitophagy, that remains sensitive to depletion or deletion of the mitochondrial deubiquitylase USP30 (Marcassa *et al*., 2018). Thus we have employed a CRISPR/Cas9 screening approach to identify further ubiquitin E3 ligases, which may regulate mitophagy.

All previous mitophagy screens have used acute mitochondrial depolarisation as a trigger for mitophagy in cell systems over-expressing Parkin (Heo *et al*, 2019; Hoshino *et al*, 2019). Here we have undertaken the technically more challenging task of screening for regulators of mitophagy in the absence of Parkin both with and without a depolarising insult. We identify two Cullin RING E3 ligase substrate adapters, VHL and FBXL4, as the most profound negative regulators under both basal and depolarising conditions. VHL is a major tumour suppressor whilst FBXL4 has been linked to early onset mitochondrial encephalomyopathy, also referred to as mitochondrial DNA depletion syndrome 13 (Bonnen *et al*, 2013; Gai *et al*, 2013; Gossage *et al*, 2015). FBXL4 is a substrate receptor of a Cullin-1 RING E3 (CRL1) ligase complex and has previously been shown to bind to SKP1, the common CRL1 adapter subunit (Tan *et al*, 2013; Winston *et al*, 1999). It encodes a mitochondrial targeting sequence and has been shown to associate with mitochondria (Bonnen *et al*., 2013; Gai *et al*., 2013). A recent study found increased levels of mitophagy in both FBXL4 KO mice and patient derived fibroblasts, although the mechanism of action remained to be elucidated (Alsina *et al*, 2020).

The connection between VHL and mitophagy is well established, operating through the control of HIF-1/2 α stability and consequent effects upon transcription of the mitophagy adaptors BNIP3 and BNIP3L/NIX, hereafter referred to as NIX (Bellot *et al*, 2009). In distinction to mitophagy adaptors utilised by the PINK1-Parkin pathway, BNIP3 and NIX integrate in the outer mitochondria membrane and then recruit the phagophore membrane via their LIR motifs in common with other selective autophagy adaptors (Hanna *et al*, 2012; Novak *et al*, 2010). We now show that FBXL4 is a suppressor of BNIP3 and NIX, but in contrast to VHL, exerts its control through the alternative mechanism of direct interaction and protein-destabilisation. Analysis of a FBXL4 mutant, associated with Mitochondrial DNA depletion syndrome 13, highlights the pathophysiological relevance of NIX as a substrate for FBXL4 and provides a molecular explanation for the aetiology of the disease. Our study indicates that NIX can be considered the master regulator of basal mitophagy and we have now unveiled a novel mechanism through which it is controlled.

## Results

### CRISPR/Cas9 E3 ligase screen for mitophagy

We chose to utilise hTERT-RPE1 cells in our search for alternative ubiquitin E3 ligases that modulate mitophagy in the absence of Parkin. We have previously established that these cells respond to depletion and deletion of the deubiquitylase USP30 by an increase in basal mitophagy, indicating that alternative ubiquitin dependent mitophagy pathways are in operation (Marcassa *et al*., 2018). We first introduced the mitochondrial matrix-targeted pH-sensitive mitophagy reporter mt-mKeima into a puromycin sensitive clone of hTERT-RPE1 cells expressing inducible Cas9 (Cas9i). This fluorophore responds to the acidic pH of the lysosomal compartment by a shift in its excitation spectrum. Hence, it reports on the delivery of mitochondrial fragments to mature autophagosomes (mitolysosomes), and provides a ratiometric readout that can be monitored by flow cytometry (Katayama *et al*, 2011). We established that our cells exhibit a measurable response to mitochondrial depolarisation using a 24 hour treatment of respiratory chain inhibitors, antimycin and oligomycin (AO). This resulted in a >2-fold increase in the population of cells captured in the high mitophagy gate (**Fig EV1A-E**). Cas9 expression could be easily detected by western blotting upon treatment with doxycycline (**Fig EV1F**).

To initiate the fluorescence based CRISPR KO screen, RPE1-Cas9i-mt-mKeima cells were incubated with a lentiviral E3 ligase sgRNA library, targeting 606 Ubiquitin E3 ligases, at a multiplicity of infection (MOI) of 0.2 and a 1000x representation. Cells were then grown in puromycin containing media to select for integration of sgRNAs prior to induction with doxycycline to initiate Cas9 expression and gene targeting (**Fig 1A**). Half of the cells were treated with AO for 24 hours to induce mitophagy, and then sorted alongside untreated cells (basal mitophagy) for both high (enhanced) and low (inhibited) mitophagy. An unsorted sample of each population was collected for reference. Genomic DNA was harvested, barcoded sgRNA libraries amplified and two independent replicates (pooled 2 by 2 from 4 independent biological experiments) were analysed by next-generation sequencing (NGS). Enriched sgRNAs were identified using the ScreenProcessing Pipeline (Horlbeck *et al*, 2016), obtaining for each comparison both a gene-level enrichment value as well as a p-value comparing the combined enrichment of all 4 sgRNAs targeting an individual E3 to that of 243 non-targeting control (NTC) sgRNA values in the same replicate (**Fig 1B, C and Fig EV2A, Dataset EV1**, 1B; average values shown). In contrast to other mitophagy screens performed in Parkin-overexpressing cells (Heo *et al*., 2019; Hoshino *et al*., 2019), we anticipated a narrow dynamic range, especially in our experiment designed to monitor basal mitophagy.

**Figure 1:**
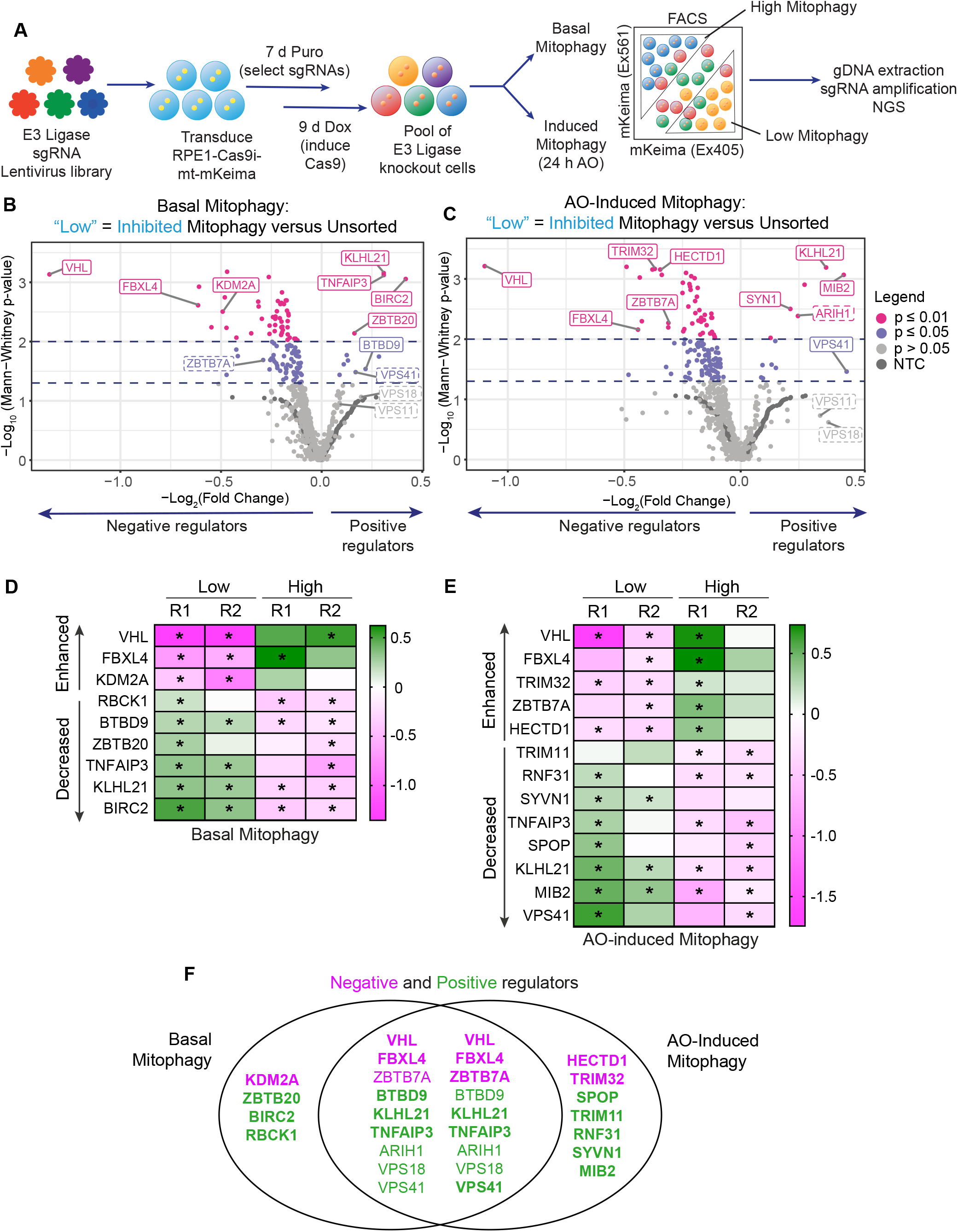
CRISPR screen targeting E3 ligases identifies major regulators of Parkin-independent mitophagy. A. Schematic depicting the CRISPR screening strategy. RPE1-Cas9i-mt-mKeima cells were transduced with a lentiviral CRISPR sgRNA library targeting 606 E3 ligases. sgRNA expressing cells were selected for 7 days with puromycin (Puro) and Cas9 expression induced with doxycycline (Dox) for 9 days. Fourteen days post transduction, half the cells were treated with AO (1 μM, 10 μM) for 24 hours, then sorted alongside untreated cells (basal mitophagy) by FACS into “high” and “low mitophagy” populations based on mt-mKeima fluorescence. Genomic DNA was extracted from sorted and unsorted reference cells, sgRNAs amplified by barcoded PCRs and samples analysed by next generation sequencing (NGS). B. Volcano plot showing the average −Log2 (Fold Change) and p-value of genes in low mitophagy vs unsorted cells in the AO-induced and basal mitophagy screen for two independent biological replicates. Statistical thresholds of two and three standard deviations from the mean are indicated by dashed lines and colour coding. Indicated are high confidence (unbroken line) and lower confidence (dashed line) candidates shown in C-E. C and D. Candidate list of positive (decreased) and negative (enhanced) regulators of Parkin-independent induced mitophagy. Heatmap showing the Log_2_(Fold Change) of genes in low/high mitophagy vs unsorted cells in the induced and basal mitophagy screen for each of two independent biological replicates. Genes with p-values <0.05 are indicated by an asterisk. E. Venn diagram showing the overlap of genes listed in C that were identified in the basal and AO-induced mitophagy screens as positive (green) and negative (magenta) regulators. Bold type indicates high confidence hits, regular type lower confidence hits.

Any sgRNAs that knock-out a positive regulator of mitophagy are expected to be enriched in the “Low” (decreased) mitophagy gates and de-enriched in the “High” (enhanced) mitophagy gates. We assembled an initial candidate list, of E3 ligases that met (de)-enrichment criteria (p<0.05; −0.2>-log2(Fold Change)>0.2) in at least one comparison in either replicate (**Dataset EV1**). A handful of E3s returned p-values <0.05 in two replicates either in the “High” or “Low” gates in basal or induced mitophagy (**Fig 1D-F**). Under mitochondrial depolarising conditions, our screen presumably represents the sum of basal and AO-induced mitophagy and it is thus expected that critical regulators of basal mitophagy are identified in both screens. Indeed, we identify 9 E3s that are common to both screens and only 4 that are unique to basal mitophagy (**Fig 1E**). Common positive regulators of basal and AO-induced mitophagy include two subunits of the HOmotypic fusion and vacuolar Protein Sorting (HOPS) complex (VPS41 and 18; VPS11 scores do not pass the thresholds set here). This provided us with confidence that our screen was identifying meaningful hits as these, and/or other HOPS complex subunits, have previously been identified in mitophagy and global autophagy screens (Heo *et al*., 2019; Hoshino *et al*., 2019; Jia & Bonifacino, 2019; Moretti *et al*, 2018; Morita *et al*, 2018; Shoemaker *et al*, 2019) and are well understood to regulate membrane fusion of autophagosomes and endosomes with lysosomes (Jiang *et al*, 2014; Takats *et al*, 2014). Five positive and two negative regulators are only detected in the AO-induced mitophagy screen.

### VHL and FBXL4 repress BNIP3 and NIX protein levels by different mechanisms

Here, we have focused our attention on the two major outliers of the basal mitophagy screen, VHL and FBXL4, for which the knock-out results in a strong increase in mitophagy (**Fig 1B-F**). VHL acts as the substrate recognition subunit of a Cullin 2-RING E3 ligase complex and is responsible for the constitutive turnover of the transcription factors HIF1α and HIF2α. Hypoxic conditions result in disengagement of HIF1α and HIF2α from VHL leading to their stabilisation and the transcription of a large number of genes, including the paralogues BNIP3 and NIX (Bellot *et al*., 2009). We wondered if FBXL4 loss may likewise promote mitophagy by upregulating BNIP3 and/or NIX.

We first adopted an orthogonal approach of depleting VHL and FBXL4, in a parental hTERT-RPE1 FlpIN cell line, with pools of siRNAs. We found a clear increase in both NIX and to a lesser extent BNIP3 protein levels in FBXL4 depleted cells, whereas BNIP3 rather than NIX was up-regulated in response to VHL depletion (**Fig 2A and B**). Importantly, in contrast to VHL knockdown, depletion of FBXL4 did not stabilise HIF1α, providing a first indication for a distinct mechanism controlling BNIP3 and NIX protein levels. Evidence that this is non-transcriptional is provided by the observation that neither BNIP3 or NIX transcript levels increase in FBXL4 depleted cells, whilst both transcripts were clearly sensitive to HIF1α depletion (**Fig EV2C**).

**Figure 2:**
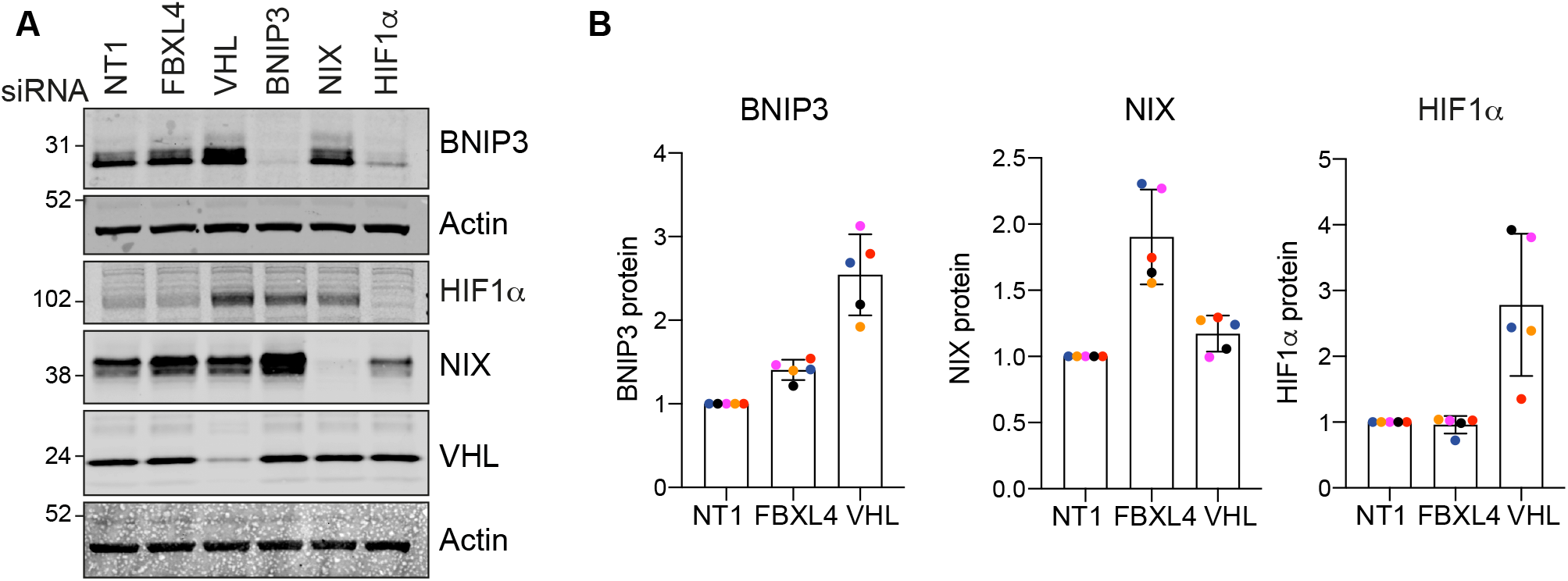
VHL and FBXL4 control expression of mitophagy adaptors BNIP3 and NIX. A. Representative western blot of hTERT-RPE1 cells transfected for 72 hours with siRNA against indicated targets or non-targeting (NT1) prior to harvesting and cell lysis. B. Quantification of BNIP3, NIX and HIF1α protein levels for data represented in A. Individual datapoint from n=5 independent colour coded experiments are shown. Error bars show standard deviation.

We next chose two sgRNAs from the E3 library mitophagy screen to generate a pair of knockout (KO) cell pools. We transduced the RPE1-Cas9i-mt-mKeima cells with the respective sgRNA encoding lentivirus, selected for stable sgRNA integration using puromycin selection and then induced Cas9 to initiate the knockout, all the while maintaining a non-induced pool of cells as a control population. Both KO pools generated with independent sgRNAs showed a clear increase in both BNIP3 (3-4x) and NIX (5-7x) protein levels and a significant increase in basal mitophagy that can be measured by both flow cytometry and live cell imaging (**Fig 3A-F**). A recent proteomic analysis comparing FBXL4 KO mice and patient derived fibroblasts reported decreased levels of a large number of mitochondrial proteins with a concomitant increase in lysosomal proteins (Alsina *et al*., 2020). Our FBXL4 KO RPE1 cells also displayed a moderate loss of the outer mitochondrial protein TOMM40 but we did not observe any significant changes in cathepsin D, one of the lysosomal enzymes highlighted in the preceding study (**Fig EV2D, E**). As expected, immunofluorescence revealed a mitochondrial localisation for this excess BNIP3 and NIX (**Fig 3C**). Importantly, we did not observe any changes in NIX or BNIP3 transcript levels, nor in HIF1α protein levels, further indicating a post-transcriptional regulatory mechanism at play (**Fig 3G, H**).

**Figure 3:**
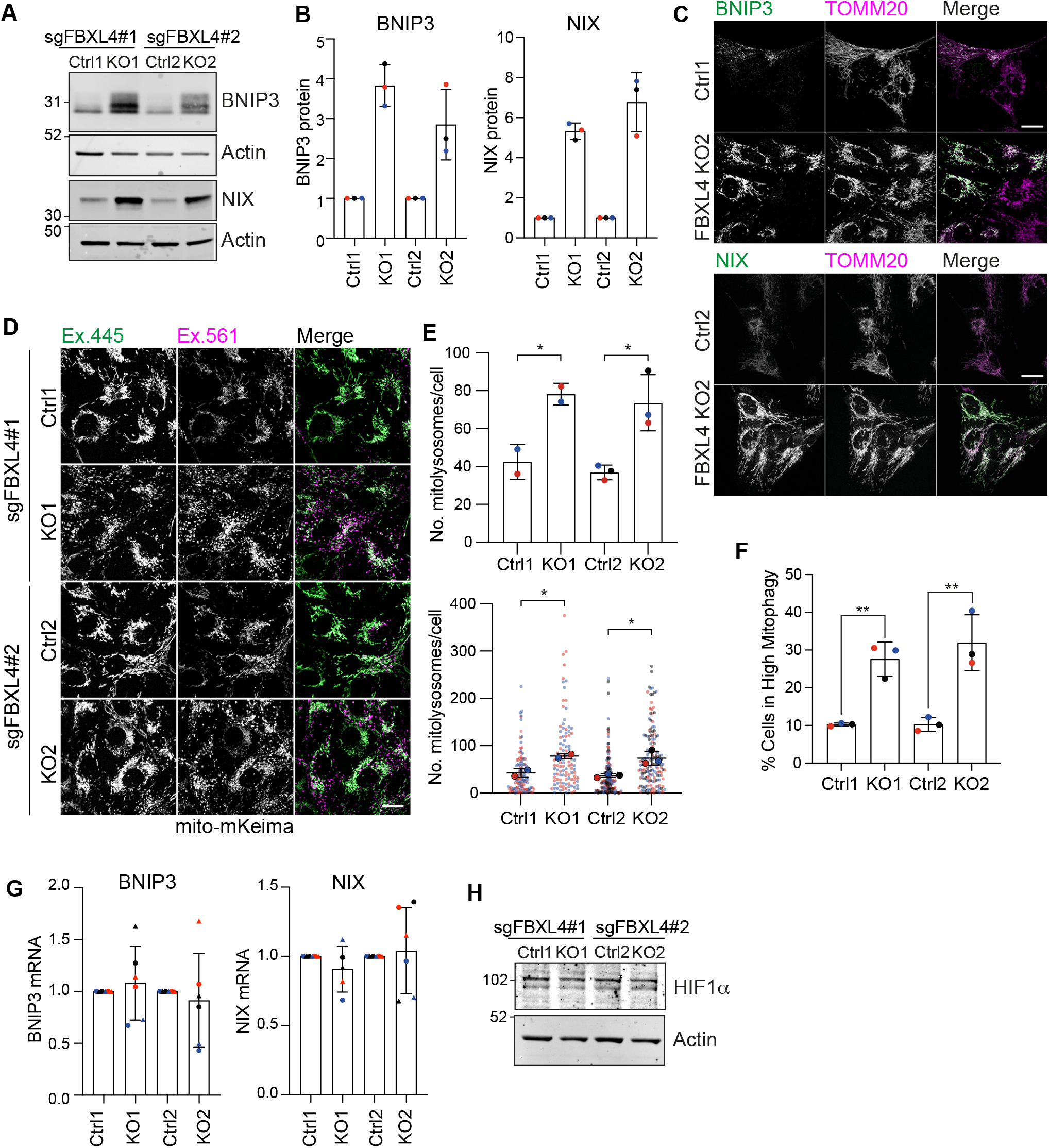
FBXL4 knockout upregulates BNIP3 and NIX and enhances basal mitophagy. A. Representative western blot of two matched pairs of control (Ctrl) and FBXL4 KO pool RPE1-Cas9i-mt-mKeima cells, expressing two distinct sgRNAs. B. Quantification of data represented in A. Individual datapoint from n=3 independent colour coded experiments are shown. Error bars show standard deviation of the fold increase of BNIP3 and NIX in each KO pool normalised to their matched control pools. C. Representative images of Ctrl and FBXL4 KO pool RPE1-Cas9i-mt-mKeima cells fixed and stained for either BNIP3 or NIX, and co-stained for TOMM20. Scale bar 20 μm. D. Representative images of control or FBXL4 KO pool RPE1-Cas9i-mt-mKeima cells. Scale bar 20 μm. E.Graphs show the quantification of mt-mkeima “red” puncta (dots) per cell. A minimum of 45 cells were analysed per condition in each experiment. Individual data from two (for control 1 (Ctrl 1) and knockout 1 (KO1)) or three (control 2 (Ctrl 2) and knockout 2 (KO2)) are shown using different colours for each experiment. Mean and standard deviation are indicated. One-way ANOVA with Tukey’s multiple comparison’s test, *P<0.05. F. Levels of mitophagy were analysed in Ctrl and FBXL4 KO RPE1-Cas9i-mt-mKeima by flow cytometry. High mitophagy gates were set using the corresponding control for each knockout. Three independent colour coded experiments are shown; one-way ANOVA and Tukey’s multiple comparison’s test, **P<0.01. G. Quantitative RT-PCR reactions of BNIP3 and NIX (normalised to Actin) were performed with cDNA derived from control and FBXL4 KO RPE1-Cas9i-mt-mKeima cells. 2 primer pairs were used per target indicated by symbol shape. Three independent colour-coded experiments are shown. Error bars show standard deviation of the fold change of BNIP3 and NIX mRNA in each KO pool normalised to their matched control pools. G. Representative western blot of control and FBXL4 KO RPE1-Cas9i-mt-mKeima cells probed for HIF1α.

### FBXL4 interacts with BNIP3 and NIX and governs their stability

We wondered whether BNIP3 and NIX could be physiologically relevant substrates of the CRL^FBXL4^. Consistent with this hypothesis, reciprocal pulldown experiments using GFP-tagged BNIP3 or NIX and Flag-tagged FBXL4 confirmed their interaction (**Fig 4A** and **B**). Disease associated mutations are largely focused in the leucine rich region which in other FBXL family proteins mediate substrate recruitment (Ballout *et al*, 2019; Bonnen *et al*., 2013; Gai *et al*., 2013). Immunofluorescence microscopy of endogenous BNIP3 and NIX showed that overexpression of wild-type FBXL4-Flag was able to suppress both BNIP3 and NIX, whereas the FBXL4 LRR mutant [R482W] was not (**Fig 4C-E**). Although only a fraction of transfected cells expressed the transgenes, a clear decrease in NIX and BNIP3 protein levels was also seen by immunoblotting in cells transfected with wild-type but not mutant FBXL4 (**Fig 4F**).

**Figure 4:**
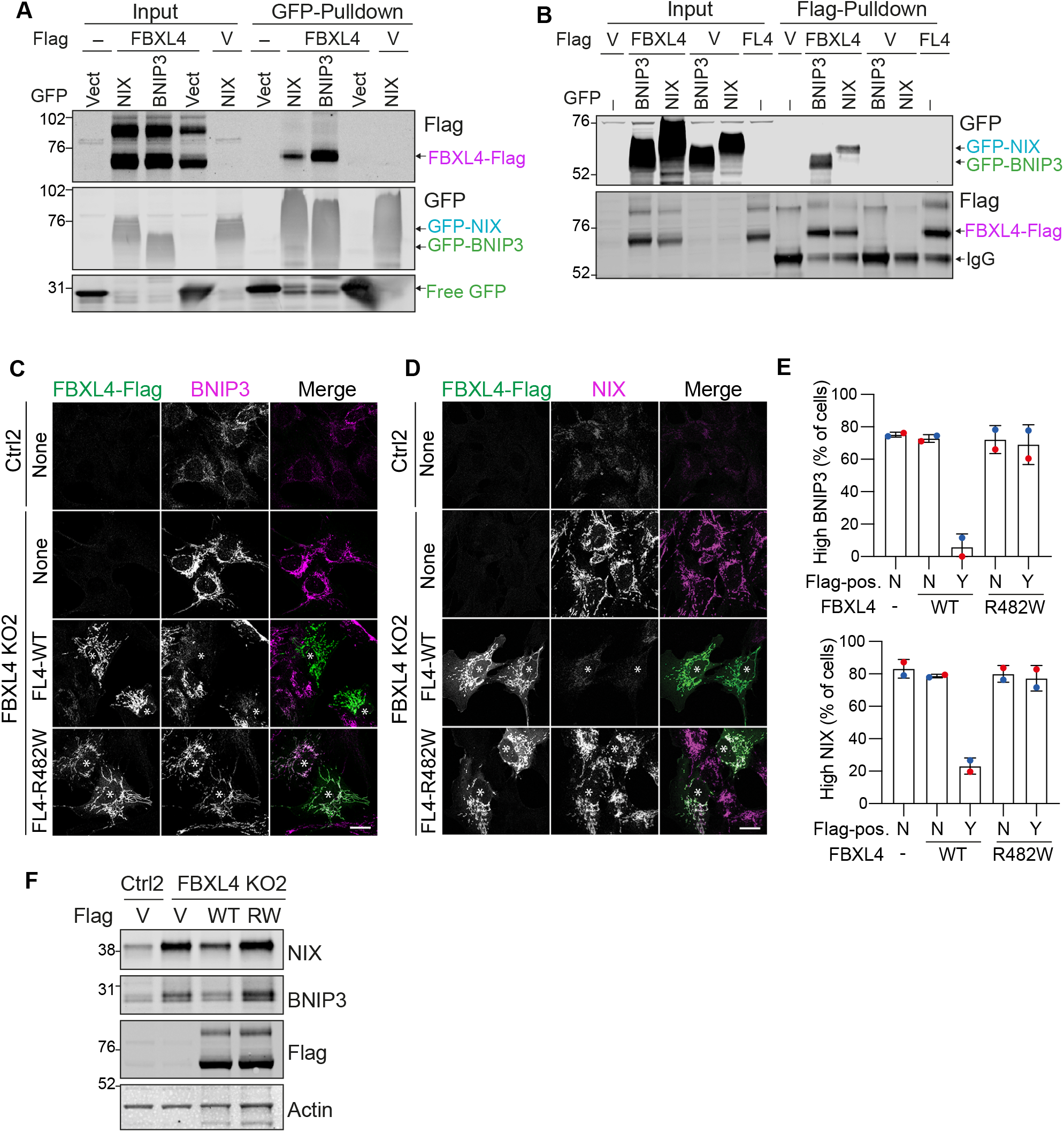
FBXL4 interacts with and destabilises BNIP3 and NIX. A and B. hTERT-RPE1 cells transfected with Flag (V) or FBXL4-Flag (FL4) were co-transfected with either GFP (Vect), GFP-BNIP3 or GFP-NIX. Lysates were subjected to immunoprecipitation (IP) with either GFP-nanobody coupled beads (A) or Flag-antibody couples agarose beads (B). IPs were probed alongside 10 μg of input sample. C and D. Representative images of Control (Ctrl) and FBXL4 KO RPE1-Cas9i-mt-mKeima cells transfected with FBXL4-Flag wild-type (WT) or FBXL4-Flag [R482W] for 24 hours, then fixed and stained for either BNIP3 or NIX. Scale bar 20 μm. E. Quantitation of the frequency of high BNIP3 and high NIX phenotypes observed in C and D. Quantitation of the frequency of high BNIP3 and high NIX phenotypes observed in C and D. A minimum of 32 Flag-positive (Flag-pos.) and Flag-negative cells were analysed per condition in each experiment. Y, Yes; N, No. Mean and standard deviation are indicated. F. Western blot of control and FBXL4 KO RPE1-Cas9i-mt-mKeima cells transfected with Flag (V), FBXL4-Flag (WT) or FBXL4-Flag [R482W] (RW) for 24 hours.

### NIX is the principal regulator of mitophagy

We reasoned that if the increased levels of BNIP3 and/or NIX are the root cause for the enhanced mitophagy in FBXL4 KO cells, then we should be able to restore low mitophagy levels by depleting these factors. Our results identify NIX rather than BNIP3 as the primary mediator of enhanced mitophagy. Depletion of NIX, but neither BNIP3, nor HIF1α, reduces basal mitophagy levels to that of the control cells (**Fig 5A, B**).

**Figure 5:**
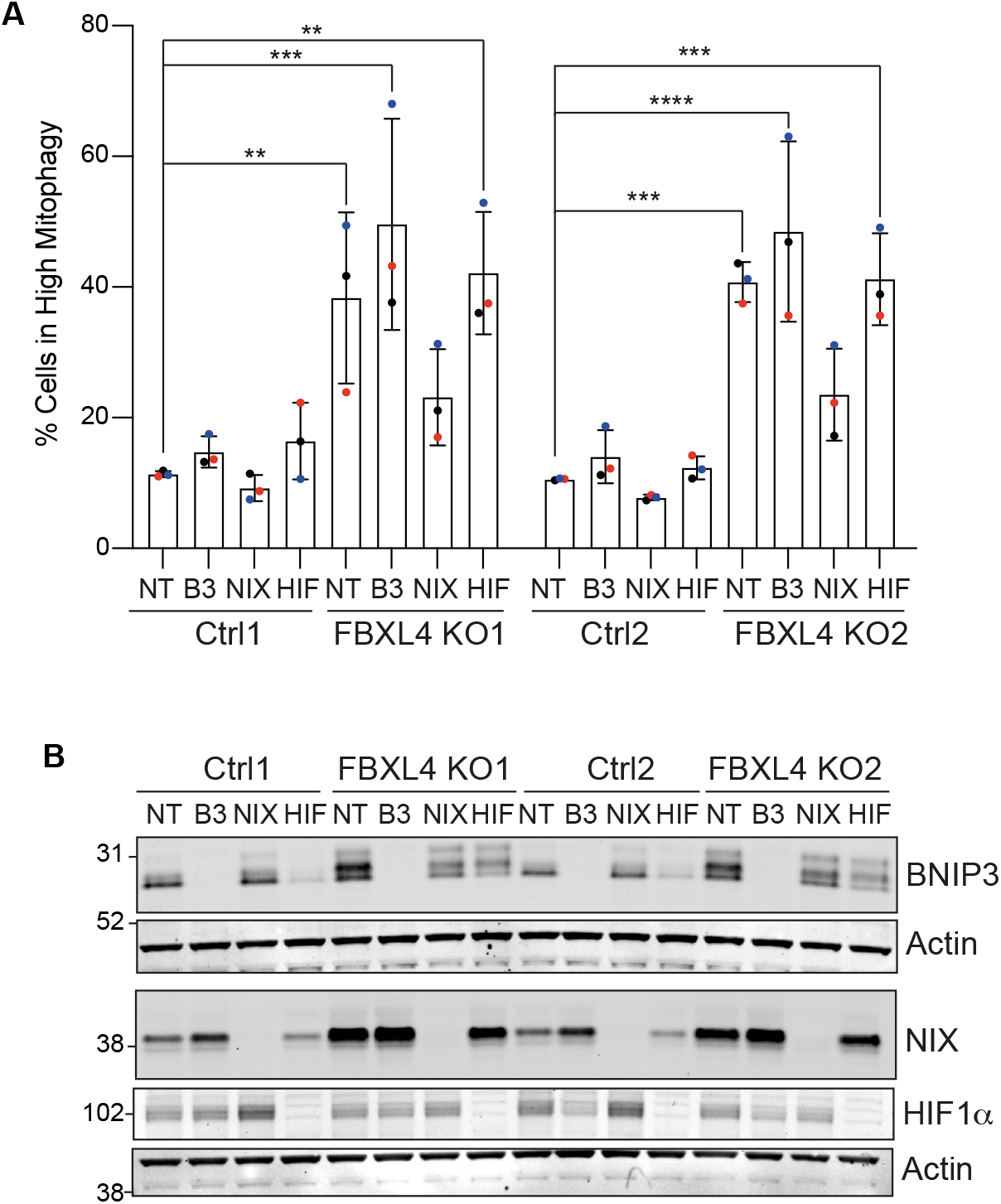
Enhancement of basal mitophagy upon FBXL4 deletion is NIX dependent. A. Levels of mitophagy were analysed in control (Ctrl) or FBXL4 KO RPE1-Cas9i-mt-mkeima cells treated with siRNA targeting the indicated proteins using flow cytometry. High mitophagy gates were set for each matched pair of control and KO cells using the Non-targeting (NT) siRNA treated Control pool sample as a reference. Data shown represent three colour-coded independent experiments. Error bars show standard deviation. One way ANOVA and Dunnett’s multiple comparison test.. B. Representative western blot of control (Ctrl) or FBXL4 KO RPE1-Cas9i-mt-mkeima cells treated with either non-targeting (NT), BNIP3 (B3), NIX of HIF1α (HIF) siRNA and probed for indicated proteins.

### Enhancement of mitophagy by MLN4924

Both VHL and FBXL4 are substrate recognition modules of CRL E3 ligase complexes, which require Neddylation for their activation. Global inactivation of CRLs can be acutely achieved with MLN4924, a highly selective compound that inhibits the Nedd8-E1 conjugating enzyme (Brownell *et al*, 2010). We surmised that MLN4924 application may provide a potent stimulus to mitophagy through up-regulation of BNIP3 and NIX.

We first treated mt-mKeima expressing hTERT-RPE1 cells for 24 hours with MLN4924 and assessed the impact on basal mitophagy by fluorescence microscopy. We observed a dramatic increase in mitophagy, which surpasses the levels elicited with a depolarising trigger (AO) over the same time period (Fig 6A). Both BNIP3 and NIX protein levels are strongly increased in the MLN4924 but not AO treated samples, whereas only AO treatment resulted in PINK1 stabilisation and an associated phosphoSer65-Ubiquitin signal (**Fig 6B-D** and **Fig EV3A**). The increase of BNIP3 and NIX proteins is seen as early as 6 hours after application of MLN4924 (**Fig 6E**). Both BNIP3 and NIX protein increase is mirrored by an increase in transcripts (**Fig EV3C**) but only BNIP3 protein levels are sensitive to HIF1α depletion (**Fig 6F, G**). The MLN4924 induced increase in basal mitophagy is sensitive to NIX-but not BNIP3 depletion alone, although a double knockdown is required for full restoration (**Fig 6H**). Depletion of HIF1α only has a marginal effect on MLN4924-induced mitophagy, which provides further support for NIX as the main mediator of this pathway (**Fig 6I**).

**Figure 6:**
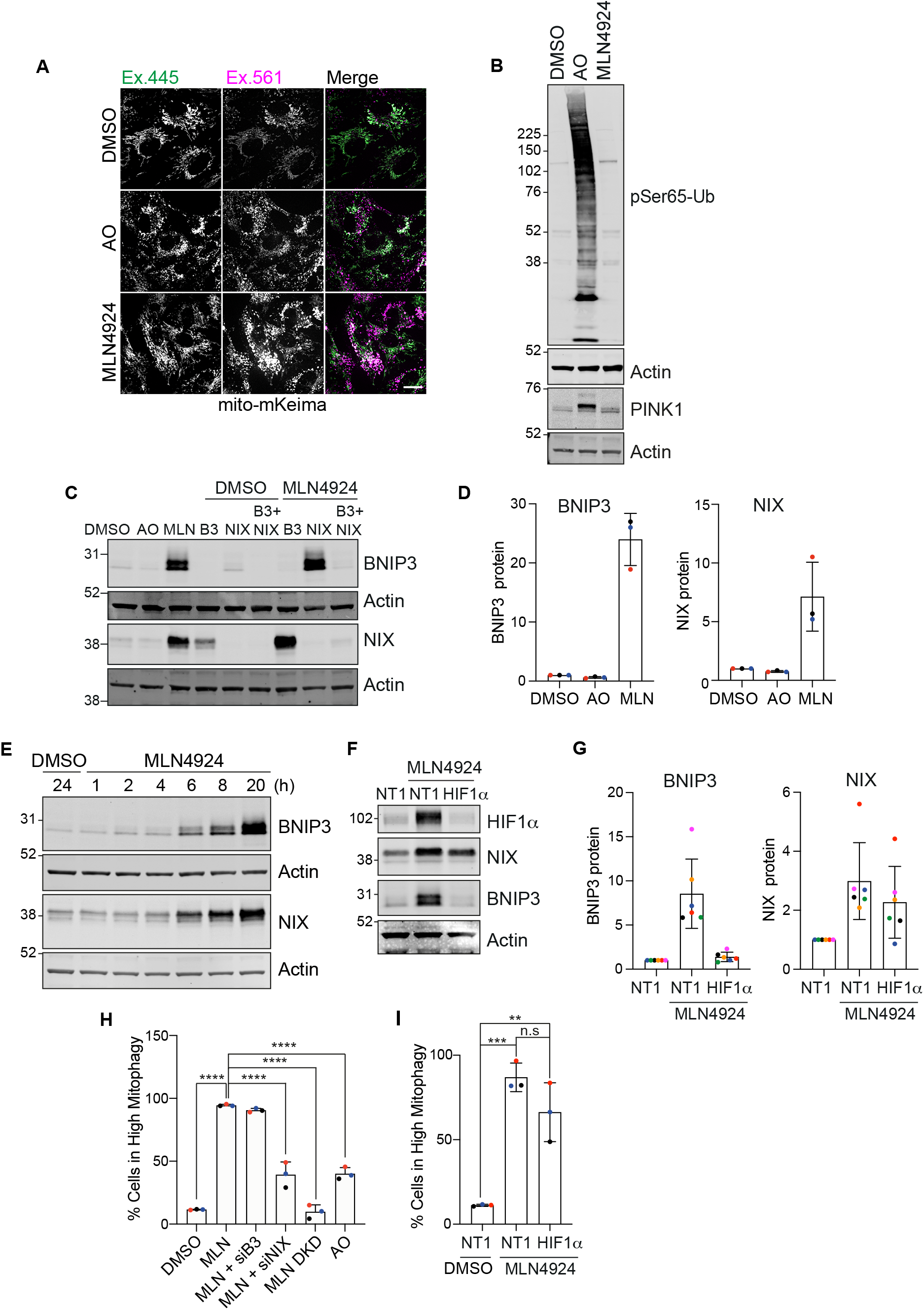
The neddylation inhibitor MLN4924 upregulates BNIP3 and NIX and enhances mitophagy. A. Representative images of RPE1-Cas9i-mt-mKeima cells treated with oligomycin (10 μM) and antimycin (1 μM) or MLN4924 (1 μM) for 24 hours. Scale bar 20 μm. B. Western blot analysis of hTERT-RPE1 cells treated with oligomycin (10 μM) and antimycin (1 μM) or MLN4924 (1 μM) for 24 hours. C. Representative western blot of hTERT-RPE1 cells transfected with siRNA targeting BNIP3 (B3), NIX or both for 72 hours. Cells were treated with either DMSO or MLN4924 (1 μM) for the last 24 hours of transfection. Untransfected cells were treated with either DMSO, oligomycin (10 μM) and antimycin (1 μM) or MLN4924 (1 μM) for the last 24 hours of the experiment. D. Quantification of data shown in C for three colour-coded independent experiments. E. Representative western blot of hTERT-RPE1 cells treated with either DMSO or MLN4924 (1 μM) for the indicated time points. F. Representative western blot of hTERT-RPE1 cells transfected with non-targeting (NT1) of HIF1α (HIF) targeting siRNA. Cells were treated with MLN4924 (1 μM) for the last 24 hours of transfection. G. Quantification of data shown in F for six colour-coded independent experiments. Error bars show standard deviation. H. Levels of mitophagy were measured in RPE1-Cas9i-mt-mkeima cells using flow cytometry. Cells were transfected with siRNA targeting BNIP3 (B3), NIX or both (DKD) for 72 hours. Cells were treated with either DMSO or MLN4924 (1 μM) for the last 24 hours of transfection. Untransfected cells were treated with either DMSO, oligomycin (10 μM) and antimycin (1 μM) or MLN4924 (1 μM) for the last 24 hours of the experiment. I. Levels of mitophagy were measured in RPE1-Cas9i-mt-mkeima cells using flow cytometry. Cells were transfected with non-targeting (NT1) or HIF1α (HIF) targeting siRNA and were treated with DMSO or MLN4924 (1 μM) for the last 24 hours of transfection. Data shown in H and I represent three colour-coded independent experiments. Error bars show standard deviation. One way ANOVA and either Tukey’s (H) or Dunnett’s (I) multiple comparison test.

By 24 hours of MLN4924 treatment, BNIP3 protein levels increase by >20x as compared to just 3-4x seen in the FBXL4 KO cells whereas the fold increase for NIX is comparable (7x) (**Fig 6D, 3B**). Importantly, in FBXL4 KO cells, NIX but not BNIP3 protein levels were refractory to MLN4924, providing strong evidence for FBXL4 as the key neddylation dependent mediator of NIX (**Fig 7A**). We noticed that the MLN4924 induced NIX protein had a shorter half-life than the basally expressed pool (**Fig EV3D**), providing an opportunity to explore the impact of FBXL4 on NIX and BNIP3 half-lives. We pretreated our isogenic cell pairs (±FBXL4) for 6 hours with MLN4924 to accumulate NIX and BNIP3 and then following MLN4924 washout, performed a cycloheximide chase experiment. Whilst BNIP3 stability appears to be marginally affected by FBXL4 KO, NIX is significantly more stable in the absence of FBXL4 (**Fig 7B-D** and **EV3E-G**). Finally we established that the turnover of this labile pool of NIX is indeed proteasome dependent by rescue with the proteasome inhibitor MG132 (**Fig 7E**).

**Figure 7:**
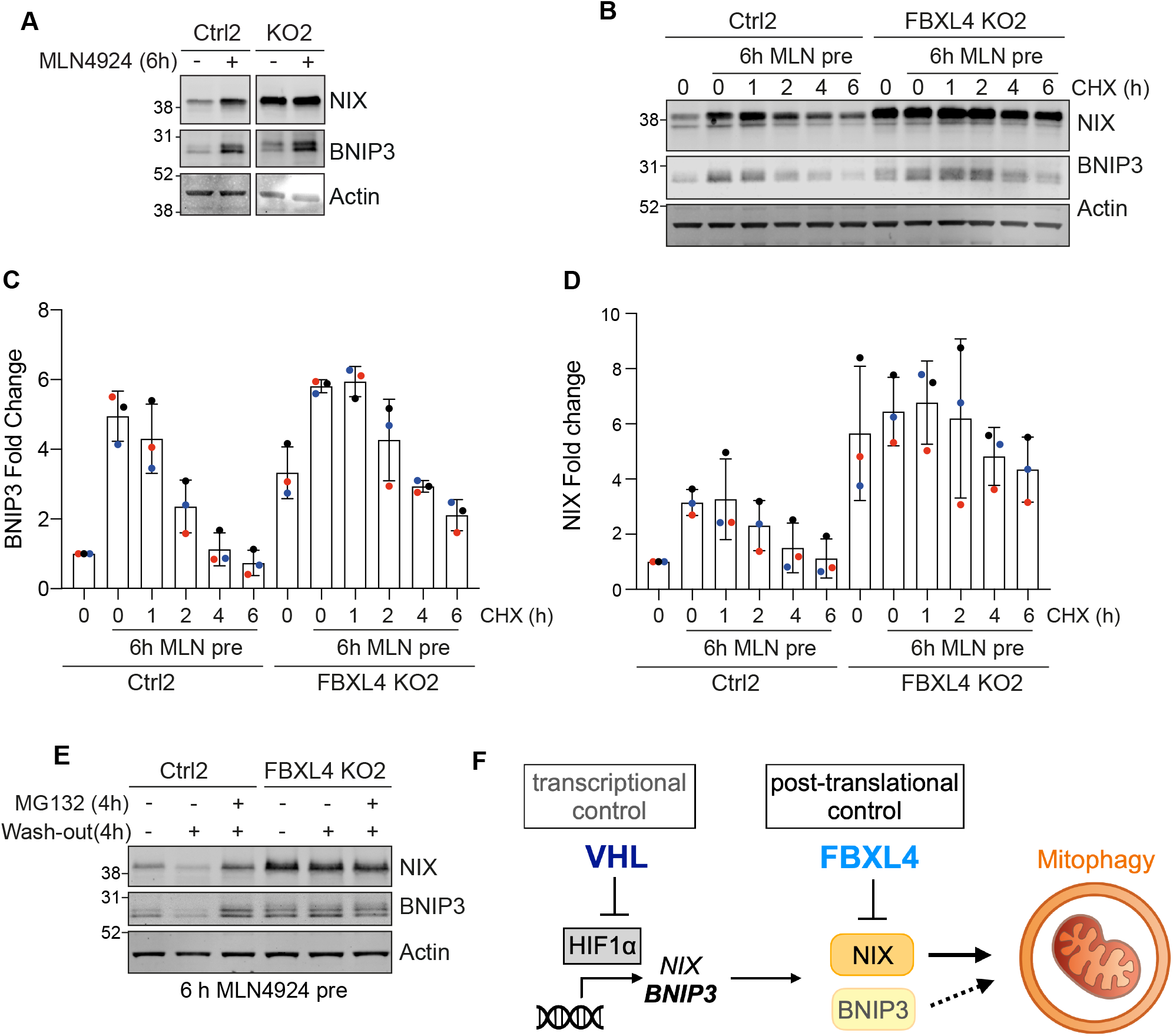
FBXL4 controls NIX and BNIP3 stability. A. Representative blot of control or FBXL4 KO RPE1-Cas9i-mt-mKeima cells treated with MLN4924 (1 μM) for 6 hours. B. Representative blot of control or FBXL4 KO RPE1-Cas9i-mt-mKeima cells pre-treated with MLN4924 (1 μM) followed by a 6-hour cycloheximide (CHX) chase (100 μg/ml). C and D. Quantification of data shown in B for three colour-coded independent experiments. Error bars show standard deviation. E. Western blot of control or FBXL4 KO RPE1-Cas9i-mt-mKeima cells pre-treated with and without MLN4924 (MLN, 1 μM) for 6 hours. MLN4924 was washed out and cells were incubated with or without MG132 (10 μM) for 4 hours before lysis. F. Schematic of proposed dual control of BNIP3 and NIX expression levels by VHL and FBXL4.

## Discussion

The critical role of mitophagy in multiple physiological settings has been established (Esteban-Martinez & Boya, 2018; Harper *et al*., 2018; Onishi *et al*, 2021; Ordureau *et al*, 2021; Schweers *et al*, 2007; Teresak *et al*, 2022; Zhao *et al*, 2020). Given the variety of contexts in which it occurs, it is not surprising that multiple mechanisms of induction and control exist. Previous mitophagy CRISPR screens have focused on the PINK1-PRKN dependent pathway which is elicited by acute mitochondrial depolarisation (Heo *et al*., 2019; Hoshino *et al*., 2019). Here we have diverged in using a cell line which does not express Parkin and have given equal emphasis to an experimental condition which does not employ depolarisation, i.e. basal mitophagy. This condition likely corresponds to the majority of mitophagy in living organisms which is PINK1-PRKN independent (Lee *et al*., 2018; McWilliams *et al*., 2018). Many of the top hits from our screen are shared between basal and depolarising conditions, suggesting that the basal mitophagy mechanisms are still extant alongside new pathways that are induced by depolarisation. However, we have also identified certain E3s, such as MIB2, that specifically promote mitophagy under depolarising conditions. These will be pursued in future studies, not least for a role in priming the PINK1-PRKN cascade.

We here identified VHL as a strong suppressor of mitophagy under both basal and depolarising conditions. The mechanism by which VHL controls NIX/BNIP3 expression, and hence mitophagy, via suppression of HIF1α is well understood. FBXL4 and KDM2A were identified as other strong suppressors of basal mitophagy. Interestingly, KDM2A is a lysine demethylase which is up-regulated by HIF1α (Batie *et al*, 2017). FBXL4 more closely corresponds with VHL, appearing as a strong suppressor hit in both conditions. We have now been able to unravel why this is so. Both VHL and FBXL4 (but not KDM2A) control BNIP3 and NIX levels but by different mechanisms (**Fig 7F**). In our system NIX, rather than BNIP3 is the crucial regulator of mitophagy. FBXL4 interacts with and regulates the stability of NIX. Proteins often display non-exponential decay rates reflective of multiple pools (McShane *et al*, 2016; Rusilowicz-Jones *et al*, 2022). Global studies of protein decay rates have revealed that supernumerary copies of proteins operating within complexes are frequently unstable. Here we find that over-production of NIX by treatment with the neddylation inhibitor MLN4924, leads to a short-lived pool that is turned over in an FBXL4 dependent fashion. One further interesting link between FBXL4 and VHL has been made in a whole genome CRISPR screen for synthetic lethality with common tumour suppressors. Loss of FBXL4 is synthetically lethal with loss of VHL, which based on our findings, we speculate is due to super-elevated levels of BNIP3 and/or NIX (Feng *et al*, 2022).

We have introduced MLN4924 as a new tool to promote basal mitophagy, predominantly through the combined inhibition of VHL and FBXL4. This will complement the widely used iron chelator deferiprone (DFP), which likely promotes mitophagy through inhibition of the iron dependent enzyme Hypoxia-inducible factor prolyl hydroxylase (Allen *et al*, 2013; Zhao *et al*., 2020). As the clinical safety profile of MLN4924 is well characterised it could be safely repurposed to stimulate mitophagy in patients who might have accumulated damaged mitochondria. Periodic stimulation of non-selective autophagy by TOR inhibitors has been shown to benefit mice bearing neurodegenerative conditions associated with aggregate formation (Menzies *et al*, 2010; Ravikumar *et al*, 2004; Rubinsztein, 2006). As a corollary one must also consider the impact of enhanced mitophagy in ongoing clinical trials for other indications.

In conclusion we have made a mechanistic link between FBXL4 and NIX in setting the levels of basal mitophagy that can explain the aetiology of early onset mitochondrial encephalomyopathy. Our screening efforts highlighted two CRLs as critical regulators of mitophagy leading us to test MLN4924 as a pharmacological mitophagy inducing agent.

We expect this will take a prominent place in the mitophagy toolbox and look forward to exploring the clinical implications.

## Materials and Methods

### Generation of a RPE1-Cas9i-PuroS cell line

To generate a Puromycin sensitive version of the CT33-hTERT-RPE1-Cas9i cell line (received as gift from Ian Cheeseman), the puromycin acetyltransferase (PAC) gene was targeted with CRISPR-Cas9. Cells were transiently transfected with pSpCas9(BB)-2A-GFP (PX458; 48138; Addgene) vector encoding a sgRNA targeting PAC (5′-TGTCGAGCCCGACGCGCGTG-3′) using Lipofectamine 2000 (Invitrogen). GFP positive cells were isolated by FACS two days after infection. Single clones were isolated by screening for susceptibility to 3 μg/ml Puromycin and indel formation was confirmed by Sanger sequencing of a PCR product isolated from genomic DNA. The selected clone, hereafter referred to as RPE1-Cas9i, contained an 18 bp deletion in the PAC gene.

### Generation of the mitophagy reporter RPE1-Cas9i-mt-mKeima cell line

A mammalian expression construct for mKeima targeted to the mitochondrial matrix via a COXVIII-derived targeting sequence (AB01-pcDNA3.1-mt-mKeima-BlastR) was transiently transfected into RPE1-Cas9i cells using Lipofectamine 2000 (Invitrogen). Transfected cells were selected with 10 μg/ml Blasticidin S HCl for 20 days and mKeima positive cells were isolated by FACS. A clonally isolated cell line expressing suitable levels of the mt-mKeima reporter was selected for all further experiments in this paper. This cell line is resistant to Hygromycin (conferred by hTERT vector pGRN145), G418/Geneticin/Neomycin (conferred by the Cas9i vector HP138-neo), Blasticidin (conferred by the vector pAB01-pcDNA3.1-mt-mKeima-BlastR), and sensitive to Puromycin.

### E3-targeting sgRNA library generation

SgRNAs targeting 606 Ubiquitin E3 ligases, included 243 non-targeting control (NTC) sequences were designed by Azimuth2 (Doench *et al*, 2016) and the top 4 picks (on-target scores>0.6) were chosen per gene, extended by 5 and 3 prime 3Cs homology and obtained as a pool from Twist Bioscience (**Dataset EV1**). The E3-targeting sgRNA library was made by 3Cs, as described previously (Diehl *et al*, 2021; Wegner *et al*, 2019; Wegner *et al*, 2020). In brief, 3Cs-DNA was generated by mixing phosphorylated pre-annealed oligonucleotides (sgRNA-encoding) with ssDNA (library template plasmid). 3Cs-DNA was purified and desalted using the GeneJET Gel Extraction Kit (Thermo Fisher) and parts of it analysed by gel electrophoresis alongside the dU-ssDNA template. The remaining 3Cs-DNA was electroporated into 10-beta electrocompetent E. coli (New England Biolabs) by using a Bio-Rad Gene Pulser and incubated over night. The final sgRNA library plasmid DNA was purified using the Maxi Plasmid DNA Prep Kit (Qiagen).

### Virus production

To produce lentiviral particles for the screen, 70% confluent Lenti-X HEK293T (Clontech) cells were transfected in 10 cm^2^ dishes with 16.5 μg E3 sgRNA library plasmid, 13.5 μg pPAX2 (12260; Addgene) and 5 μg pMD2G (12259; Addgene) using Lipofectamine 2000 according to the manufacturer’s instructions. The media containing lentiviral particles was harvested 48 hours later, centrifuged at 500 g for 5 minutes and small aliquots were snap frozen in liquid nitrogen and stored at –80 °C. The viral titre was determined by serial dilution and polybrene mediated transduction of RPE1-Cas9i cells followed by selection in 5 μg/ml Puromycin.

### CRISPR library screen

RPE1-Cas9i-mt-mKeima cells (75% confluency) were transduced on Day 0 with a lentiviral CRISPR library containing 2667 unique sgRNA encoding viruses, targeting 606 E3 ligases with 4 sgRNAs per gene plus 243 non-targeting controls (NTC) using polybrene (8 μg/ml). To achieve >1000x library coverage at a multiplicity of infection of 0.2, a total of 13.5 × 10^6^ cells were transduced for each replicate and maintained throughout the screen. 24 hours after transduction, Puromycin selection (5 μg/ml) was initiated and maintained for 7 days. On day 3, Cas9 expression was induced with Doxycycline (1 μg/ml) and maintained for 10 days. On day 14, “induced mitophagy” was initiated by treatment with Antimycin A (1 μM) and Oligomycin A (10 μM), while “basal mitophagy” was allowed to proceed in untreated cells. Approximately 24 hours later, cells were harvested separately for FACS. A fraction of 2.7 × 10^6^ cells were snap frozen as a cell pellet on liquid nitrogen as an unsorted reference sample for both induced and basal mitophagy.

### Fluorescence-activated cell sorting and flow cytometry

For the CRISPR/Cas9 mitophagy screen, cells were harvested by trypsinisation, washed, resuspended in FACS buffer (1% FBS/PBS) and stored on ice. Cells were sorted using a FACSAria III (BD Biosciences). Mitolysosomal versus mitochondrial mt-mKeima fluorescence was measured using dual-excitation ratiometric pH measurements at 488 nm (pH 7) and 552 nm (pH 4) lasers with 695/40 nm and 610/20 nm emission filters, respectively. The top (high mitophagy) and bottom (low mitophagy) 20-30% of cells were collected in FACS buffer, washed with PBS and snap frozen in liquid nitrogen. Four biologically independent experiments were sorted for each sample.

Samples for flow cytometry were prepared as above. Initial characterisation of RPE1-Cas9i-mt-mKeima by flow cytometry was done using a LSR Fortessa (BD Biosciences) using 405 nm (pH 7) and 561 nm (pH 4) lasers with 610/20 nm emission filters for the LSR Fortessa. Data were analysed using Flowjo (v 10). High mitophagy gates were made by taking the top 10-15% of the control sample and applied to the other samples. Post screen validation by flow cytometry was carried out using either a Bio-Rad ZE5 Cell analyser or a Sony MA900 cell sorter, using dual-excitation ratiometric pH measurements. pH 7 and pH 4 measurements were taken using 488 nm and 552 nm lasers respectively and 610/20 nm emission filters. Data was analysed using Flowjo (v 10). High mitophagy gates were made by taking the top 10-15% of the control sample and applied to the other samples.

### DNA extraction, library preparation, and next generation sequencing

Genomic DNA from sorted cells was purified using a Purelink Genomic DNA Kit (Invitrogen) and eluted in 50 μl nuclease free water according to the manufacturer’s instructions. Up to 5 μg genomic DNA was typically obtained from this process. Library amplification PCRs were performed using NEBNext Ultra II Q5 Master Mix (New England Biolabs) on all the extracted genomic DNA to attach Illumina sequencing adaptors and barcodes. A universal pool of 8 forward primers were used for each sample along with a unique reverse primer containing a sample-specific barcode. The PCR cycling conditions used were 98°C for 30 seconds, 22 cycles of 98°C for 10 seconds, 63°C for 30 seconds and 65°C for 45 seconds, followed by a final extension of 65°C for 5 minutes. The library PCRs from two independent biological experiments were pooled for each of the two replicate samples submitted for NGS to ensure each NGS sample was derived from between 2.4 – 5.4 × 10^6^ cells, i.e. a ∼900-2000x coverage of the E3 sgRNA plasmid library. Samples were purified using two rounds of Ampure-XP beads (Beckman Coulter) at a ratio of 0.8x according to the manufacturers instructions. 1 μl purified product was run on a 3% agarose gel to ensure the primers were removed by purification and the amplicons were present at 375 bp. Samples were sequenced on a NovaSeq 6000 using Novaseq S4 PE150 chemistry (Novogene).

### Screen analysis

Raw sequencing reads were trimmed to 35 bp (containing unique sample barcodes) with BBTools using BBDuk (BBMap – Bushnell B. – sourceforge.net/projects/bbmap/) and untrimmed reads were removed. The ScreenProcessing pipeline (available at https://github.com/mhorlbeck/ScreenProcessing; (Horlbeck *et al*., 2016)) was used to count the number of uniquely mapped sgRNAs for each sample and to determine the enrichment of sgRNAs between samples. Briefly, the number of uniquely mapped sgRNAs for each sample was counted and sgRNAs with less than 50 reads were removed from the analysis. For each comparison, both a gene-level enrichment value (Log_2_(Fold Change) calculated from the average of all four sgRNAs for each gene), as well as a Mann-Whitney p-value, reporting the probability that the distribution of enrichment values for all sgRNAs targeting a particular gene is different to the distribution of all 243 non-targeting control (NTC) gRNA values in the same replicate was generated (Bassik *et al*, 2013; Kampmann *et al*, 2013) (**Dataset EV1**). Plots were generated in R using ggplot2 and modified in Adobe Illustrator for style.

### Cell culture, transfection, siRNA interference

RPE1 cells were cultured in Dulbecco’s modified Eagle’s medium DMEM/F12 (hTERT-RPE1) supplemented with 10% FBS and 1% non-essential amino acids, and Lenti-X HEK293T cells were cultured in Dulbecco’s modified Eagle’s medium DMEM (Gibco) supplemented with 10% FBS. For siRNA experiments, cells were treated with 40 nM of non-targeting (NT1) or target-specific siRNA oligonucleotides (Dharmacon On-Target Plus), using Lipofectamine RNAi-MAX (Invitrogen, 13778030) according to manufacturer’s instructions. Medium was exchanged after 6 h and cells harvested 72 h after transfection. For plasmid transfections, Lipofectamine 2000 (Invitrogen, 11668019), Lipofectamine 3000 (Invitrogen, L3000001) or Genejuice (Merck millipore, 70967) was used according to the manufacturers protocol, unless otherwise stated. Transfection reactions were carried out for 16-24 hours. For cycloheximide assay, cells were treated for the indicated time points with 100 μg/ml cycloheximide.

### siRNA and plasmids

Sequences of siRNA used in this manuscripts were as follows: FBXL4 (ON-TARGET Plus pool: 5’-GCAGUUGUGUCAUGAUUGA-3’, 5’-GGACAUAUUAGGAACAAGA-3’, 5’-GGAAUGGACAGUCUUAACA-3’, 5’-UUAGAAUUCUCGCUUGUUC-3’), VHL (ON-TARGET Plus pool: 5’-CCGUAUGGCUCAACUUCGA-3’, 5’-AGGCAGGCGUCGAAGAGUA-3’, 5’-GCUCUACGAAGAUCUGGAA-3’, 5’-GGAGCGCAUUGCACAUCAA-3’), BNIP3 (ON-TARGET Plus pool: 5’-UCGCAGACACCACAAGAUA-3’, 5’-GAACUGCACUUCAGCAAUA-3’, 5’-GGAAAGAAGUUGAAAGCAU-3’, 5’-ACACGAGCGUCAUGAAGAA-3’), NIX (ON-TARGET Plus pool: 5’-GACCAUAGCUCUCAGUCAG-3’, 5’-CAACAACAACUGCGAGGAA-3’, 5’-GAAGGAAGUCGAGGCUUUG-3’, 5’-GAGAAUUGUUUCAGAGUUA-3’), HIF1α (ON-TARGET Plus Pool: 5’-GAACAAAUACAUGGGAUUA-3’, 5’-AGAAUGAAGUGUACCCUAA-3’, 5’-GAUGGAAGCACUAGACAAA-3’, 5’-CAAGUAGCCUCUUUGACAA-3’). The plasmids pCDNA5 FRT TO FBXL4-3xFlag, pCDNA FRT TO FBXL4 [R482W]-3xFlag, pCDNA5 FRT TO GFP-BNIP3, pCDNA5 FRT TO GFP-NIX were all purchased from Dundee MRC PPU. The PAC sgRNA (5′-TGTCGAGCCCGACGCGCGTG-3′) (Lambrus et al., 2016) was cloned into the pSpCas9(BB)-2A-GFP (PX458; 48138; Addgene) vector while the FBXL4 sgRNA-1 (5’-TTGGTCAGAGAGACCTACGA-3’) and FBXL4 sgRNA-2 (5’-TGGACTACCTCTGCATTGAG-3’) plasmids were cloned into the pLenti-sgRNA (71409; Addgene) vector. All sgRNAs were cloned using BsmBI restriction sites. The neomycin resistance gene was removed from pcDNA3.1mito-mKeima (Katayama *et al*., 2011) with XmaI/BstBI (NEB) and the vector was treated with Mung Bean Nuclease and Alkaline Phosphatase Calf Intestinal (New England Biolabs). The blasticidin resistance gene was isolated from a plasmid pKM808 using XhoI/SalI (New England Biolabs) and treated with a Quick Blunting Kit (New England Biolabs) and the insert and vector were ligated using Quick Ligase (New England Biolabs) to generate pAB01-pcDNA3.1mito-mKeima-BlastR.

### RNA isolation and qRT-PCR

Total RNA was isolated from RPE1 cells or RPE1 control and FBXL4 KO pools using Qiagen RNA extraction kit (74106). cDNA was generated from 1 μg RNA using RevertAir H Minus reverse transcription (Thermo Scientific, 11541515) with RNasin (Promega, N251S), oligo (dT) 15 primer (Promega, C1101) and PCR nucleotide mix (Promega, U144B). Quantitative PCRs were performed in triplicate using primers against BNIP3 (Pair 1; 5’-CCTCAGCATGAGGAACACGA-3’; 5’-AAAAGGTGCTGGTGGAGGTT-3’; Pair 2; 5’-CCTTCCATCTCTGCTGCTCTC-3’; 5’-TGGAGGTTGTCAGACGCCTT-3’), Nix (Pair 1; 5’-AGGAAAATGAGCAGTCTCTGCC-3’; 5’-TGGAGGATGAGGATGGTACG-3’; Pair 2; 5’-TGTCGTCCCACCTAGTCGAG-3’; 5’-GCTGTTCATGGGTAGCTCCA-3’) or Actin (5’-CACCTTCTACAATGAGCTGCGTGTG-3’; 5’-ATAGCACAGCCTGGATAGCAACGTAC-3’). Primer and cDNA reactions were run with iTaq Mastermix (Bio-Rad, 172-5171) in a Biorad CFX Connect real-time system. The mean cycle threshold (Ct) values were normalised to Actin (ΔCt = Ct target – Ct Actin), raised to the exponent of 2^-ΔCt^ and normalised to the respective control cell line to generate 2^-ΔΔCt^.

### Antibodies and reagents

Antibodies and other reagents are as follows: anti-Actin (66009, 1:10000, Proteintech), anti-BNIP3 (ab109362 1:1000 WB, 1:100 IF, abcam), anti-NIX (12396, 1:1000 WB, 1:250 IF, Cell signalling technology), anti-HIF1α (NB100-134, 1:1000, Novus Bio techne), anti-VHL (68547, 1:1000, Cell signalling technology), anti-TOMM20 (612278, 1:1000 WB, 1:500 IF, BD transduction), anti-TOMM40 (NBP2-38289, 1:1000, Novus Bio techne), anti-VDAC1 (ab14734, 1:1000, abcam), anti-Cathepsin D (219361, 1:2000, Cal Biochem), anti-Flag (F3165, 1:1000WB; 1:250 IF, Sigma-Aldrich), anti-Flag (F7425, 1:1000, Sigma-Aldrich), anti-GFP (In house, 1:1000), anti-Phospho-Ubiquitin (Ser65) (62802, 1:1000, Cell Signalling technology), anti-Cas9 (ab191468, 1:1000, abcam), anti-PINK1 (6946, 1:1000, Cell signalling technology), anti-LC3 (5F10, 1:200, Nanotools), anti-p62 (610833, 1:1000, BD Transduction), MLN4924 (C-1231, Chemgood), MG132 (Sigma-Aldrich), cycloheximide (Sigma-Aldrich) oligomycin A (75351; Sigma-Aldrich), and antimycin A (A8674; Sigma-Aldrich).

### Preparation of cell lysates and western blot analysis

Cultured cells were lysed with RIPA buffer (150 mM NaCl, 1% Sodium deoxycholate, 10 mM Tris-Cl pH 7.5, 0.1% SDS, 1% Triton X-100) supplemented with MPI (mammalian protease inhibitor) cocktail (P8340; Sigma-Aldrich) and Phosstop (04906837001; Roche). Proteins were resolved using SDS-PAGE (Invitrogen NuPage gel 4–12%) transferred to nitrocellulose membrane (10600002; Amersham), blocked in 5% milk (Marvel) or 5% BSA (41-10-410; First Link) in TBS (20 mM Tris–Cl, pH 7.6, and 150 mM NaCl) supplemented with Tween-20 (10485733; Thermo Fisher Scientific), and probed with primary antibodies overnight. Visualisation and quantification of Western blots were performed using IRdye 800CW (goat 926-32214, mouse 926-32212 and rabbit 926-32213), and 680LT (goat 926-68024, mouse 926-68022 and rabbit 926-68023) coupled secondary antibodies and an Odyssey infrared scanner (LI-COR Biosciences).

### Coimmunoprecipitation

Cells were lysed in NP40 buffer (0.5% NP40, 25 mM Tris pH 7.5, 100 mM NaCl, 50 mM NaF) supplemented with MPI, Phosstop and CAA (2-Chloroacetamide, 50 mM), 16 hours after transfection. Lysates (250-400 μg protein) were then incubated with 10 μl of either GFP nano-trap beads or Flag affinity gel. For GFP nano-trap pulldowns, incubation was for 1 hour before beads were washed in IP wash buffer (0.1% NP40, 25 mM Tris pH 7.5, NaCl 100 mM, NaF 50 mM). Bound proteins were eluted in sample buffer (62.5 mM Tris pH 6.8, 3% SDS, 10% glycerol, 3.2% β-mercaptoethanol). For Flag pulldowns, incubation was for 2 hours before beads were washed with TBS buffer (10 mM Tris pH 7.5, 100 mM NaCl). Bound proteins were eluted in Flag peptide (Sigma, F4799, 150 ng/μl) for 30 minutes. Eluted proteins were diluted in sample buffer and analysed by western blot.

### Immunofluorescence and live cell imaging

Cells were fixed in 4% paraformaldehyde in PBS and permeabilised with 0.2% Triton X-100 in PBS. Coverslips were blocked in goat serum, then cells stained with primary antibodies. Proteins were visualised using AlexaFluor-488 or AlexaFluor-647 coupled secondary antibodies. Fixed coverslips were imaged using either a Zeiss LSM800 or LSM900 Airyscan (63x oil, acquisition software Zen blue). Images were processed using Adobe Photoshop (v 22.1.1) and Fiji (v 2.1.0) software. All images were acquired sequentially. For live-cell imaging of mt-mKeima, cells were seeded onto IBIDI μ-Dishes (81156; IBIDI) 2 days prior to imaging with a 3i Marianas spinning disk confocal microscope (63x oil objective, NA 1.4, Photometrics Evolve EMCCD camera, acquisition software Slide Book 3i v3.0). Images were acquired sequentially (445 nm excitation, 617/73 nm emission; 561 nm excitation, 617/73 nm emission). Analysis of basal mitophagy levels was performed using the “mitoQC Counter” plug-in in Fiji (v 2.1.0) software, as previously described (Montava-Garriga *et al*, 2020), using the following parameters: radius for smoothing images = 1.25, ratio threshold = 0.8, and red channel threshold = mean + 1 SD. Mitophagy analysis was performed for 2-3 independent experiments with minimum 47 cells per condition.

### Statistic analysis

Bar graphs indicate mean and standard deviation. Statistical significance was determined with an unpaired t-test (**Fig EV3C**) or one-way ANOVA with either Dunnett’s (**Fig 5A, 6H, EV2C**) or Tukey’s (**Fig 3B, 3E, 3F, 6I**) multiple comparisons tests using GraphPad Prism 9.. P-values are represented as *\*P*<0.05, *\*\*P*<0.01, *\*\*\*P*<0.001, *\*\*\*\*P*<0.0001.

## Acknowledgments

We thank Jonathon Pines (ICR London) and Iain Cheeseman (MIT) for generously providing hTERT-RPE1-FlpIN and CT33-hTERT-RPE1-Cas9i cell lines, respectively, for this project. We also thank Francesco Barone for help in evaluating and selecting the single cell m-mtKeima expressing cell clone, Jin-Rui (Amos) Liang for help and advice with the screen processing pipeline, and Robert Taylor (Newcastle, UK) for useful discussions. Initial observations of MLN4924 induced mitophagy were made by Elena Marcassa and Jane Jardine (University of Nantes, France) whilst working in the Liverpool laboratory. This work was supported by a Parkinson’s UK Grant (G-1902) and the Deutsche Forschungsgemeinschaft (DFG, German Research Foundation; Project ID 259130777 – SFB 1177). KRM is supported by a studentship from the MRC Discovery Medicine North (DiMeN) Doctoral Training Partnership (MR/N013840/1). MJC is the recipient of a Royal Society Industry Fellowship (INF\R2\212031). AJB is the recipient of funding from the European Union’s Horizon 2020 research and innovation programme under the Marie Sklodowska-Curie grant agreement No 897783.

## Figure legends

**Expanded View Figure 1:**
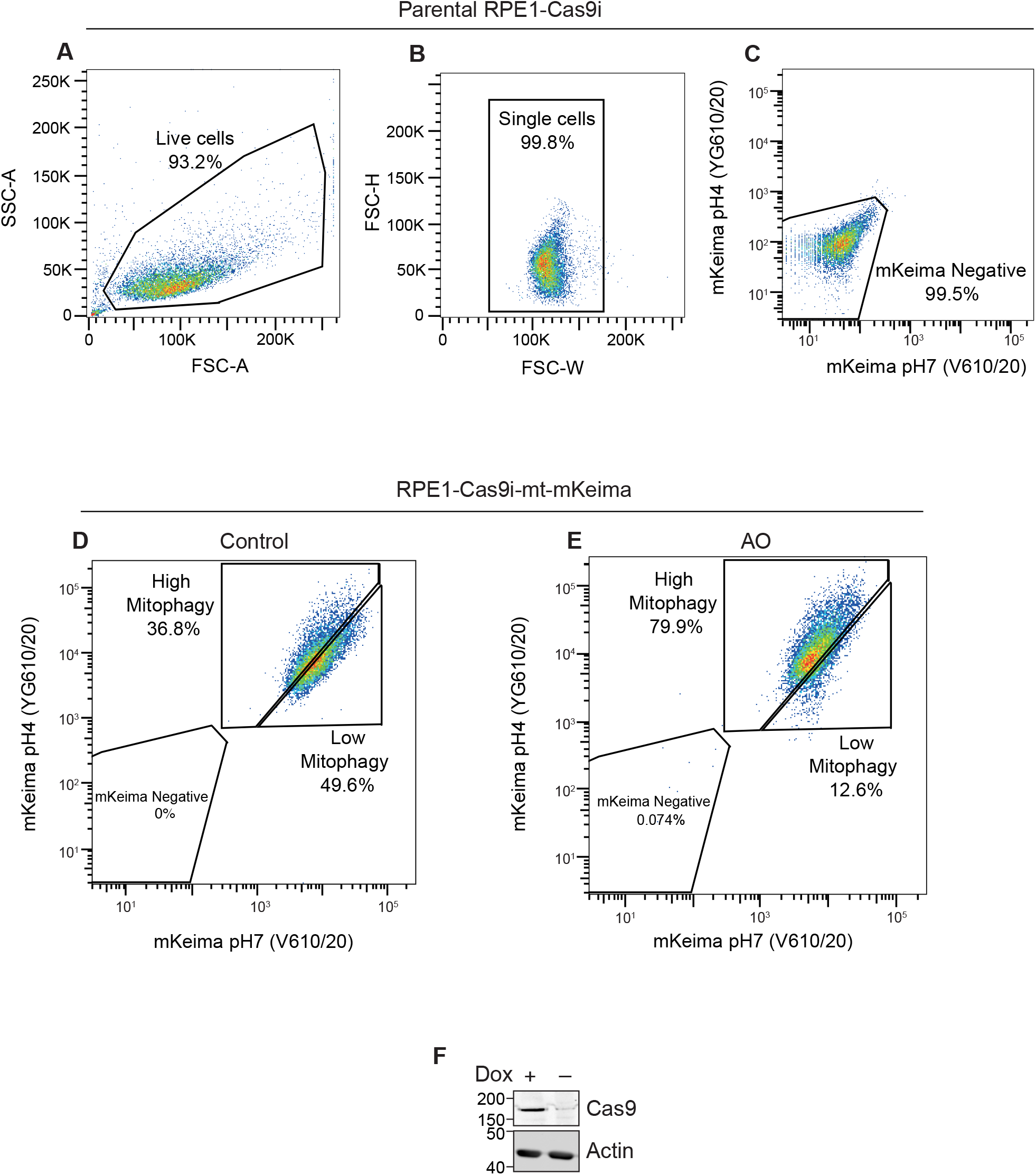
Characterisation of RPE1-Cas9i cells stably expressing mt-mKeima. A-C. Parental RPE1-Cas9i cells were used to set the gates for flow cytometry for live (A) and single cells (B), and to exclude non-fluorescent cells (C). D and E. RPE1-Cas9i-mt-mKeima cells were treated with DMSO (D) or Antimycin A and Oligomycin A (AO; 1 and 10 μM respectively) (E), for 24 hours. An increase in the proportion of cells undergoing mitophagy (high mitophagy population) is observed in AO treated cells. F. Western blot of RPE1-Cas9i-mt-mKeima for Cas9 following 4 days of Doxycycline (Dox, 1 μg/ml) induction.

**Expanded View Figure 2:**
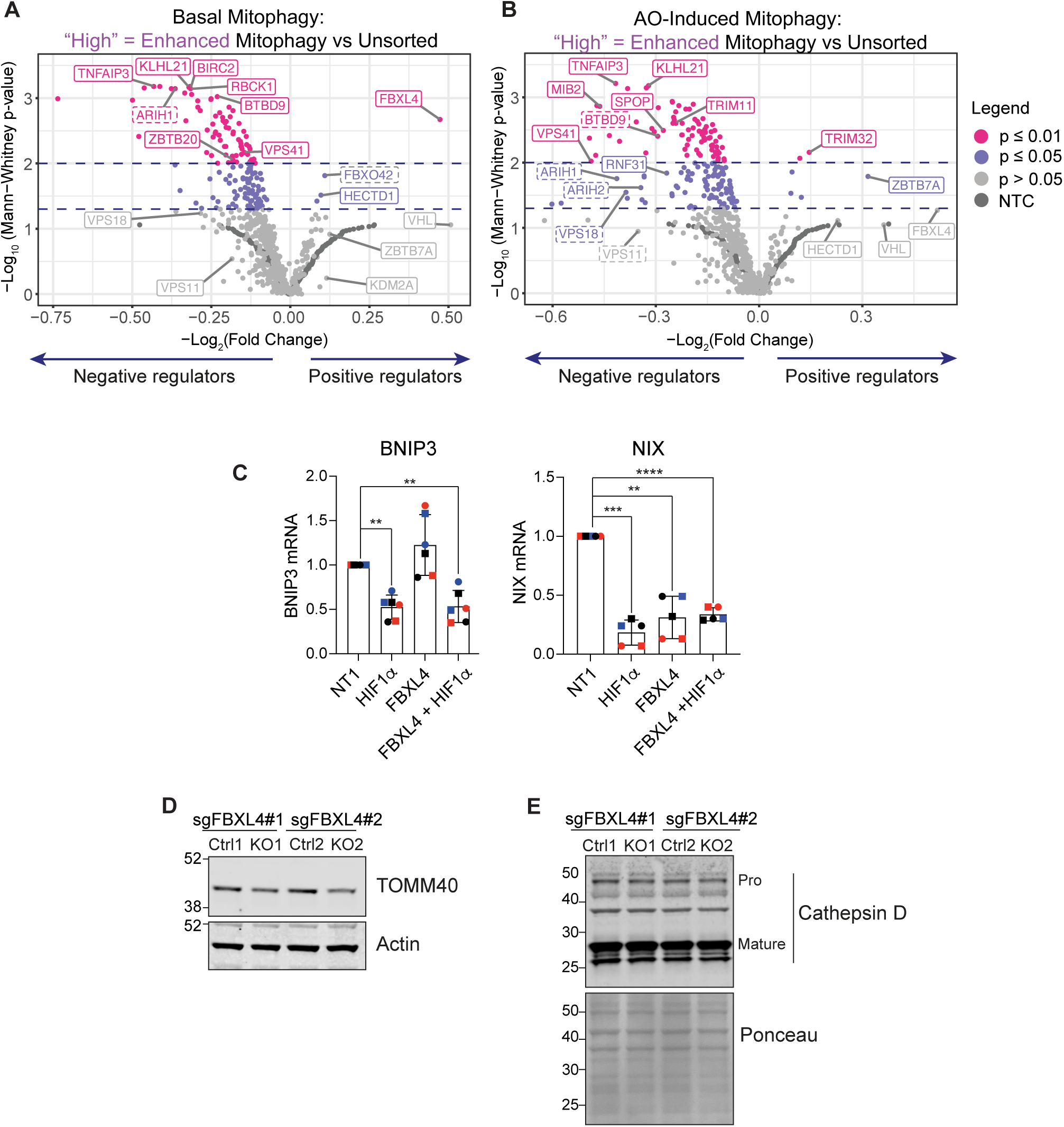
Supplementary data for Figures 1, 2 and 3. A and B. Volcano plots showing the average −Log_2_(Fold Change) and Log_10_(p-value) of genes in high mitophagy vs unsorted cells in the induced and basal mitophagy screen for two independent biological replicates. Statistical thresholds of two and three standard deviations from the mean are indicated by dashed lines and colour coding. Indicated are high confidence (unbroken line) and lower confidence (dashed line) candidates shown in Fig 1 C-E. C. Quantitative RT-PCR reactions of BNIP3 or NIX (normalised to Actin) were performed with cDNA derived from hTERT-RPE1 cells transfected with non-targeting (NT1) or siRNA targeting FBXL4 or HIF1α (HIF). Two primer pairs were used per target indicated by different symbol shapes. Shown is the fold change normalised to the NT1 control samples for three independent colour coded experiments. Error bars show standard deviation; one-way ANOVA and Dunnett’s multiple comparison’s test, **P<0.01. D. and E. Western blot of lysates from matched control (Ctrl) and FBXL4 knockout RPE1-Cas9i-mt-mkeima cell pools. Bands corresponding to Mature and Pro-Cathepsin D (Pro) are indicated.

**Expanded View Figure 3:**
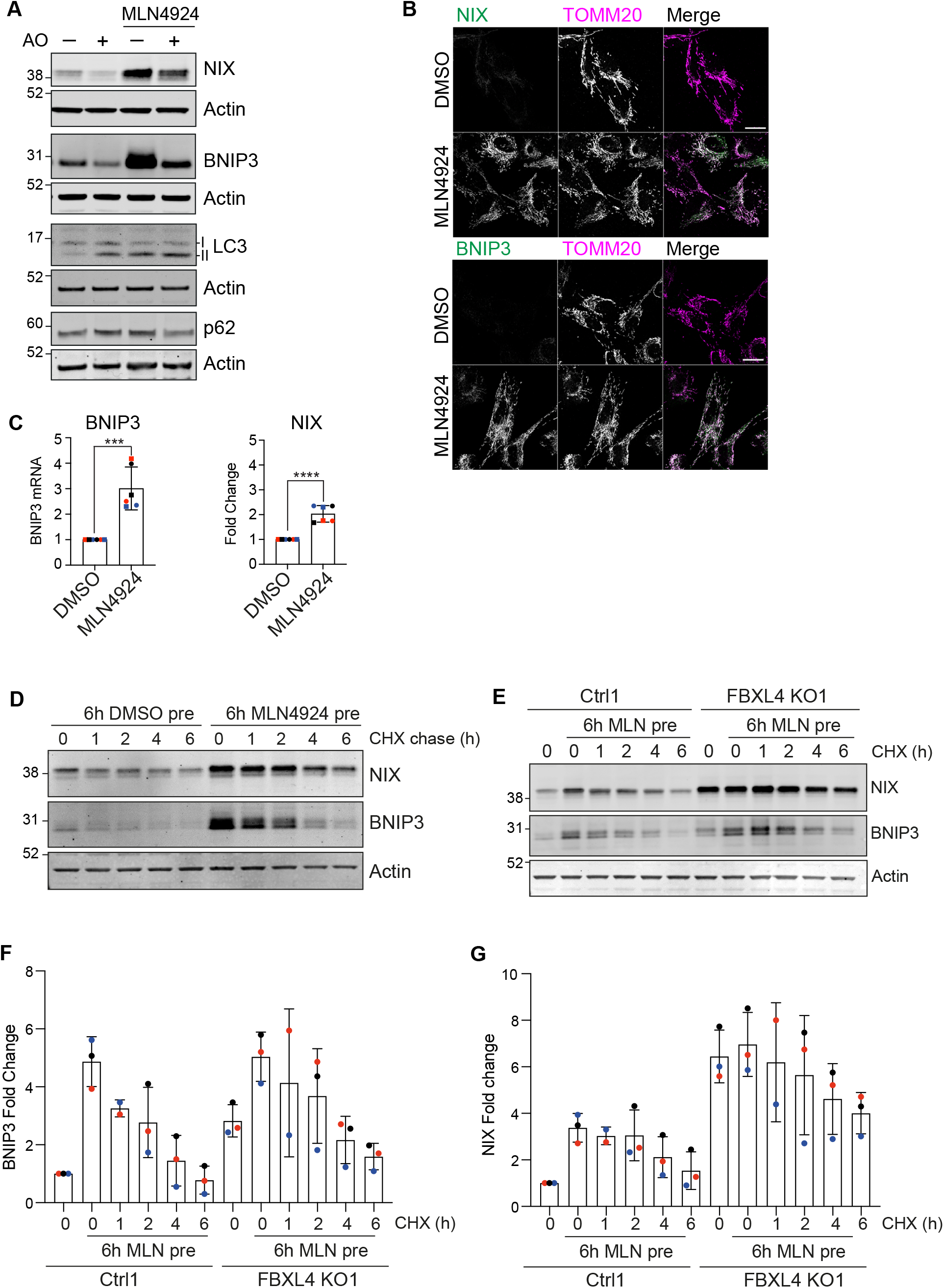
Accessory data indicating FBXL4 influences NIX and BNIP3 stability. A. Representative western blot of hTERT-RPE1 cells treated with DMSO, Antimycin (1 μM) and oligomycin (10 μM) (AO) or MLN4924 (1 μM) for 24 hours. B. Representative images of hTERT-RPE1 cells treated with DMSO or MLN4924 (1 μM) for 24 hours, fixed and stained for either BNIP3 or NIX and for TOMM20. Scale bar 20 μm. C. Quantitative RT-PCR reactions of BNIP3 or NIX (normalised to Actin) were performed with cDNA derived from hTERT-RPE1 cells treated with DMSO or MLN4924 (1 μM) for 24 hours. Two primer pairs were used per target indicated by different symbol shapes. Shown is the fold change normalised to the DMSO control sample for three independent colour coded experiments. Error bars show standard deviation. Unpaired t-test. D. Representative western blot of hTERT-RPE1 cells pretreated with DMSO or MLN4924 (1 μM) for 6 hours, followed by a 6-hour cycloheximide chase (100 μg/ml). E. Representative blot of control or FBXL4 KO RPE1-Cas9i-mt-mkeima cells pre-treated with MLN4924 (1 μM) followed by a 6-hour cycloheximide (CHX) chase (100 μg/ml). F And G. Quantitation of data shown in E for three independent colour-coded experiments, normalised to the DMSO-treated matched control line sample. Error bars show standard deviation.

### Expanded View Dataset 1

Tables showing sgRNA used to create the E3 ligase CRISPR library as well as non-targeting control sgRNAs. Tables show Log_2_(fold change) values for targeted genes as well as Mann Whitney p values generated using the Screen Processing pipeline.

